# Detoxifying and depolymerizing microorganisms reveal intertwined guild collaborations in the gut microbiome of a generalist macro-algivorous fish

**DOI:** 10.1101/2025.11.04.686673

**Authors:** Alvaro M. Plominsky, Aaron Oliver, Carlos Henriquez-Castillo, Sheila Podell, Jeremiah J. Minich, Simona Augyte, Jennica Lowell-Hawkins, Neil A. Sims, Eric E. Allen

## Abstract

The biotransformation of macroalgal biomass represents a major catabolic challenge due to its structurally diverse polysaccharides and inhibitory polyphenols. Unlike terrestrial lignocellulosic substrates, macroalgae polysaccharides contain multiple monomer types, branching patterns, and sulfation states. Additionally, macroalgae polyphenols have been shown to inhibit both microbial growth and their catalytic enzymes. While herbivorous fishes have evolved specialized gut microbiota to process these substrates, the enzymatic pathways remain poorly characterized, with few experimentally validated polysaccharide utilization loci or biochemically defined marine sulfatases, and limited understanding of polyphenol degradation. Here, we developed *in vitro* microcosms, based on the gut microbiome of the generalist macro-algivorous fish *Kyphosus cinerascens*, to temporally resolve the activity of the microbial guilds involved in macroalgal polysaccharide and polyphenol transformation. First, parallel cDNA/DNA amplicon sequencing were employed to distinguish the natural active fraction from transient gut microbiome taxa. Four media combinations were able to propagate between 96% to 99% of the active hindgut microbial families, reproducing the cooperative degradation dynamics observed *in vivo*. Metagenomic and metatranscriptomic profiling of these four optimized *in vitro* microcosms served as models to assess the stepwise functional successions occurring in the natural gut microbiome. Early Gammaproteobacteria expressed enzymes linked to polyphenol detoxification and alginate degradation, followed by Bacillota, Bacteroidota, and Verrucomicrobiota guilds targeting more recalcitrant sulfated polysaccharides and polyphenols. Together, these results identified temporal and taxonomic coordination as key features of macroalgal biomass deconstruction, providing an experimentally tractable model for discovering novel carbohydrate-active enzymes and elucidating poorly understood pathways of marine polyphenol degradation.

**IMPORTANCE:** Seaweed represents a source of sustainable biomass for various applications, but scalable industrial methods struggle to break down seaweed biomass into intermediate products due to the complexity of its constituents. Fish of the genus *Kyphosus* feed on different seaweed types by leveraging gastrointestinal bacteria to neutralize inhibitory polyphenols and convert their polysaccharides into simple sugars. This study identifies microbial groups that are transcriptionally active in natural fish hindgut microbiomes to propagate these active microbial communities *in vitro*. This enabled assessing how distinct microbial guilds act in succession to transform complex polysaccharides and polyphenols. Notably, this is the first study to assess the biotransformation capacities of macroalgal polyphenols by complex *in vitro* hindgut microbiomes of a generalist herbivorous fish. These findings advance our ecological understanding of cooperative degradation in marine gut symbioses and establish a tractable platform for discovering new enzymes and pathways with potential applications in algal biomass utilization.

## INTRODUCTION

The biotransformation of macroalgae biomass represents a major catabolic challenge due to the complexity of its polysaccharides (Arnosti et al., 2021) with diverse monomer types, branching patterns, sulfation states, and the presence of polyphenols capable of inhibiting enzymes and microbial growth (Cotas et al. 2020); (Khan et al. 2022); (Lavecchia et al. 2024). These features distinguish macroalgal biomass from terrestrial lignocellulosic substrates and demand highly coordinated enzymatic strategies for effective biotransformation.

Marine herbivorous fishes have evolved specialized gut microbiota that enable them to neutralize the chemical defenses and digest the structurally complex polysaccharides of macroalgae. The carbohydrate-active enzymes (CAZymes) and sulfatases that mediate this process are distributed across multiple microbial taxa. Individual genomes lack the complete enzymatic repertoire to fully depolymerize macroalgae sulfated polysaccharides by themselves, underscoring the need for cooperative extracellular degradation (Podell et al. 2023); (Oliver et al. 2024); (Facimoto et al. 2024). Such coordination is often encoded within polysaccharide utilization loci (PULs), which combine hydrolytic enzymes, sulfatases, transporters, and regulators. Yet, experimentally validated PULs for marine polysaccharides remain scarce, with less than 4% of known clusters linked to specific substrates (Ausland et al. 2021). Likewise, the diversity of sulfatase enzymes required to cleave polysaccharide sulfate esters is only beginning to be appreciated, with just a handful of marine carbohydrate-active sulfatases biochemically characterized to date (Hettle, Vickers, and Boraston 2022).

These gaps limit our capacity to predict pathway completeness *in silico*, especially for the most structurally complex substrates. Moreover the presence of polyphenols adds another layer of complexity. While they function as secondary metabolites that protect macroalgae from herbivory and oxidative stress, their microbial degradation pathways remain poorly understood (Cotas et al. 2020). Databases such as CAMPER (McGivern et al. 2024) are only beginning to capture the diversity of human gut microbiome genes involved in polyphenol metabolism (B. Zheng et al. 2022); (Balzerani et al. 2022), leaving open questions about how these compounds shape microbial succession and enzymatic activity in the gut environment of marine hosts. Investigating these interactions is essential, as the detoxification of polyphenols likely constrains the accessibility of underlying polysaccharides and mediates broader community dynamics during macroalgal digestion (Cotas et al. 2020).

*Kyphosus* fish provide a natural model system to address these knowledge gaps. These fishes consume a wide variety of red, green, and brown macroalgae (Wu et al. 2022; Choat, Clements, and Robbins 2002); (Sparagon et al. 2022) and play key ecological roles in coral reef systems by regulating algal cover and mediating coral–algal competition (Dell et al. 2020). Previous studies have taxonomically characterized (Sparagon et al. 2022); (Pisaniello et al. 2023) and laid the genomic foundation for the capacities of the *Kyphosus* gut microbiota to biotransform a variety of macroalgae sources (Facimoto et al. 2023); (Podell et al. 2023); (Facimoto et al. 2024). These metagenomic studies, assessing principally the distal end (hindgut) section of the host intestine, have revealed that the *Kyphosus* microbiomes are enriched in novel CAZyme and sulfatase families not found in terrestrial herbivores (Podell et al. 2023), and that many of these enzymes are phylogenetically distinct from known sequences in public databases (Oliver et al. 2024). Moreover, recent *in vitro* cultivation efforts have offered a glimpse into the advantages of using a reproducible experimental setting to study these microbiomes (Oliver et al. 2024). Overall, these studies have shown that degradation of macroalgae by the the *Kyphosus* microbiomes is carried out by taxonomically diverse groups, most prominently members of the Bacteroidota and Bacillota phylum, that partition enzymatic functions in a stepwise division of labor (Facimoto et al. 2023); (Podell et al. 2023); (Facimoto et al. 2024); (Oliver et al. 2024).

Together, these findings highlight both the ecological and biotechnological importance of *Kyphosus* gut symbionts. Ecologically, they illustrate how gut microbes underpin the ability of herbivorous fishes to utilize diverse macroalgal diets and thereby shape reef ecosystems. Biotechnologically, they offer a largely untapped resource for novel enzymes and microbial communities with potential applications in bioenergy, bioremediation, and seaweed-based aquaculture feeds (Oliver et al., 2024). Yet, while genomic and compositional insights have expanded rapidly, relatively few studies have integrated longitudinal transcriptional data or experimentally propagated whole communities to capture active enzymatic processes under controlled conditions.

This study identified the transcriptionally active members of the *Kyphosus cinerascens* hindgut microbiota, a fish species with a generalist herbivorous feeding ecology encompassing green, brown, and red macroalgae. To propagate these active communities under controlled conditions, we screened different culture media and established *in vitro* microcosms (Fig. S1), from which we collected paired 16S rRNA amplicon, shotgun metagenomic, and shotgun metatranscriptomic sequencing data across temporal stages of algal biomass degradation. Together, this data provides unprecedented resolution into the stepwise, collaborative processes underlying macroalgal degradation in herbivorous fish and establishes a reproducible experimental platform for exploring microbial succession, enzyme functionality, and community-level strategies for polysaccharide and polyphenol biotransformation.

## RESULTS AND DISCUSSION

### Amplicon-based profiling of natural and microcosm enrichments

Active *K. cinerascens* hindgut microbiota potentially involved in macroalgae biotransformations were identified by comparing cDNA- and DNA-based 16S rRNA gene amplicon datasets, using a high-throughput “Fast-Extract” DNA extraction procedure. This approach distinguished between active taxa and transient microbiota members that were ingested along with dietary items but became inactive/dead during their passage through the GI tract. Both datasets were generated from the same six *K. cinerascens* hindgut samples that served as source inocula for all microcosms (Fig. S1). At the phylum level, amplicon sequence variants (ASVs) assigned to Cyanobacteriota (potentially representing chloroplasts from the ingested macroalgae), and Epsilonbacteraeota/Campylobacterota varied greatly between DNA and cDNA abundances, with active (cDNA) fractions representing less than half of total DNA levels (Fig. 1A). The phyla Bacillota, Deinococcota/Deinococcus-Thermus, Elusimicrobiota/*Ca*. Termite Group 1, and Mycoplasmatota differences were less dramatic, but still statistically significant, with higher representation of their ASVs in the DNA versus cDNA fractions. Spirochaetota had between 20% higher representation in the cDNA (active) compared to the DNA (transient) fractions (Fig. 1A).

**FIG 1.**
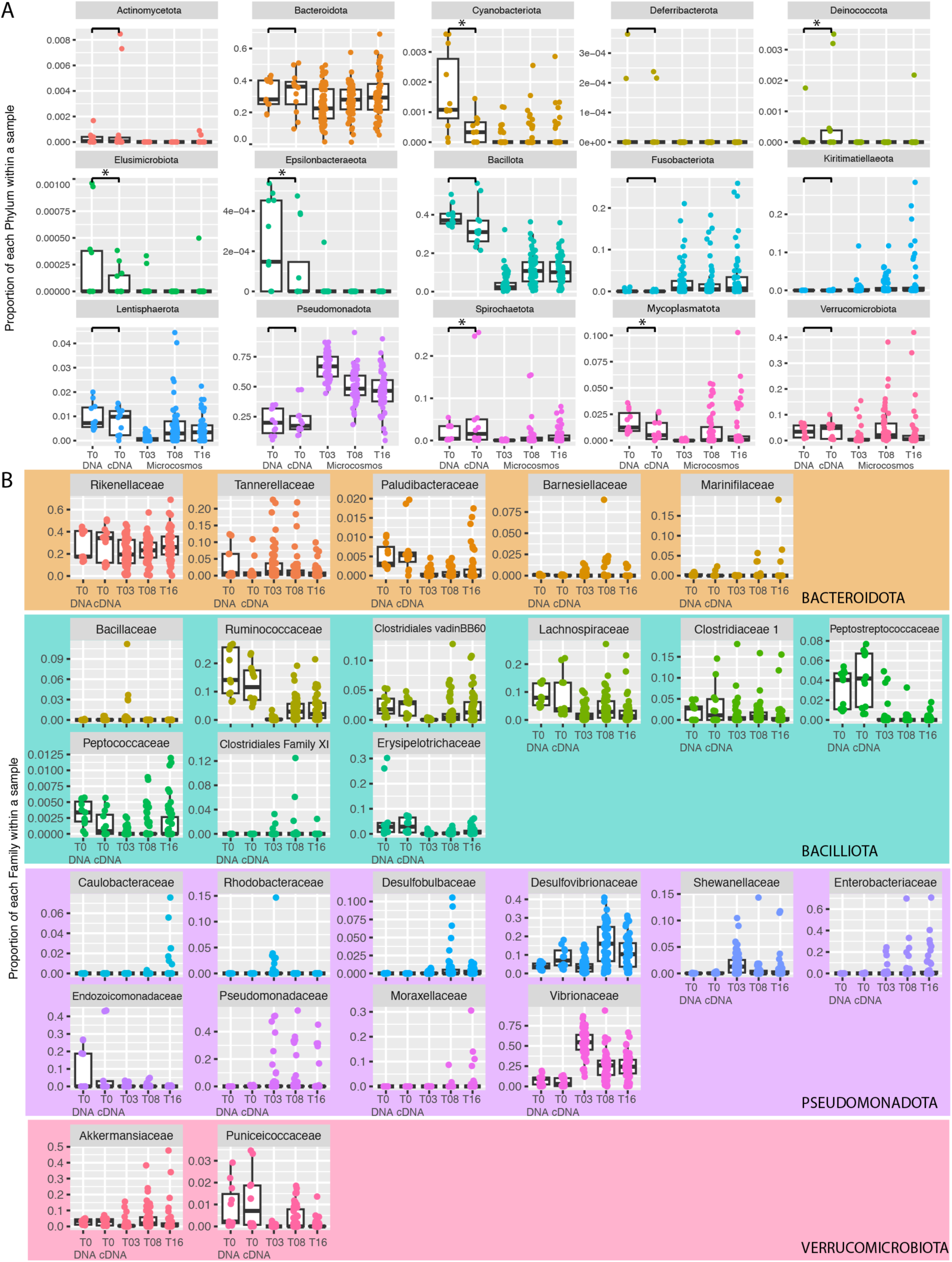
Proportional abundance of natural and propagated microbial communities of the *Kyphosus cineratus* hindgut microbiome at A) phylum and B) family level taxonomic resolution. X axis positions in each graph denote total DNA and cDNA fractions at the time of inoculation (T0), followed by results at 3, 8, and 16 days post-inoculation (T03, T08, and T16). Significant differences (p < 0.05) of mean abundances between natural *K. cinerascens* DNA and cDNA samples for each taxonomic group based on Tukey’s test are marked with an asterisk Family level abundances for Fusobacteria, Kiritimatiellaeota, Lentisphaerota, and Spirochaetota are not shown because nearly all of their taxonomically assignable reads were classified as belonging to a single family (*i.e*., Fusobacteriaceae, Kiritimatiellaceae, Victivallaceae, and Brevinemataceae, respectively). Taxa shown represent only those present at levels greater than 5×10^-4^ % of all taxonomically assignable reads in at least one sample.

ASVs assigned to microbial families that were highly represented in the cDNA (active) fractions were also found in the microcosms (Fig. 1B), although ASVs assigned to phylum Bacillota were generally less abundant in the microcosms than initial inocula. In contrast, *in vitro* cultivation conditions seem to have favored some Pseudomonadota families that became overrepresented at distinct, sporadic time-points (Fig. 1B). These results were consistent with a previous study showing high expression levels of genes for polysaccharide biotransformation were in Bacteroidota and Bacilliota representatives from the hindgut of another *Kyphosus* species (Facimoto et al. 2023).

The taxonomic compositions of microcosms across all media combinations over 16 days of cultivation were compared to determine the extent to which native *K. cinerascens* hindgut microbiota were being propagated *in vitro*. Four different media combinations (microcosms number 30, 36, 70, and 78; Fig. S1 and Table S2) propagated 96%-99% of all ASVs assignable at the family level (Fig. S4). Although breakaway diversity estimates indicated that less than one-third of this diversity was replicated all the way down to individual ASV levels (Fig. S3),22 of the 25 microbial families comprising more than 5×10^-4^% of the total ASVs in at least one sample were effectively propagated in culture (Fig. S4). One caveat is that several Pseudomonadota families that were not detected in the cDNA fraction of natural *K. cinerascens* hindgut microbiomes (*i.e*., Caulobacteraceae, Rhodobacteraceae, and Desulfobulbaceae) were detected within the total DNA fraction, perhaps being reactivated during *in vitro* cultivation (Fig. S4).

The Association Networks framework and subsequent Redundancy Analysis of ASV co-occurrence patterns were used to corroborate which microcosm media best propagated the natural hindgut taxa. A network of samples was generated based on the similarity of their ASV co-occurrence patterns (Pearson correlation coefficients ≥ 0.7) to identify strongly associated pairs. Then their modularity was assessed through Markov Cluster assignments of highly connected sample clusters that shared similar community structures. In this network, microcosm #78 formed the most connected cluster with the active (cDNA fraction) *K. cinerascens* hindgut microbiomes of fish 3, 4, and 6 (Cluster 1; Fig. S5A), indicating a strong compositional similarity. The correlation network guided the subsequent Redundancy Analysis, which tested how specific media components contributed to the propagation of these complex microbial assemblages. The fact that microcosm #78, with such a complex media composition, was the best in propagating the *K. cinerascens* hindgut active microbiome is consistent with the results showing that a combination of many of the media components, rather than a single component by itself, was required to enable the propagation of the natural hindgut community majority (Fig. S5B).

The media selected for use in detailed microcosm analyses (Fig. S1) contained several different combinations of macroalgae. Microcosm 30 contained (brown) *Turbinaria* spp. plus various (red) Coralline representatives. Microcosm 36, also included (green) *Ulva* spp., Microcosms 75 and 78 contained the most complex macroalgae mixtures, encompassing (brown) *Turbinaria* spp., *Halidrys* spp., and *Dictyota* spp., various (red) Coralline representatives, and (green) *Ulva* spp. The main distinction between the media of microcosms 75 and 78 was the addition of propionic acid, casein, and butyric acid in the latter (Fig. S1).

### Metagenomic and metatranscriptomic profiling of natural and microcosm enrichments

The homogeneity of the combined *K. cinerascens* natural hindgut inoculum used to make all microcosms was corroborated by simultaneous metagenomic (Table S5) and metatranscriptomic analysis (Fig. 2B and C), which also confirmed the propagation of active microbial members from the natural *K. cinerascens* hindgut microbiota in the four selected microcosms (Fig. 2). However, as expected when comparing metagenomic and 16S rRNA amplicon data derived from different nucleic acid extraction procedures (*i.e.*, Fast-Extract versus a commercial Zymo kit), there were differences in the proportional abundances of certain groups. Such was the case for Alphaproteobacteria, which were underrepresented at the 16S rRNA gene level (Fig. 2A) but were prevalent across all time points of these four selected microcosms at both the metagenomic and metatranscriptomic levels (Fig. 2B and C). In contrast, the representation of Spirochaetota and Verrucomicrobiota was higher at the 16S rRNA gene level (Fig. 2A) compared to their representation assessed through metagenomic and metatranscriptomic approaches (Fig. 2B and C). The metagenomic data generated from the natural fish hindguts, the natural hindgut inoculum, and all three time points of the selected microcosms enabled the binning of 1,153 metagenome-assembled genomes (MAGs; Table S6). The propagation of populations from the main taxonomic groups comprising the natural *K. cinerascens* hindgut microbiome was further confirmed by the taxonomic redundancy observed between the MAGs generated from the source inoculum and the selected microcosms (Fig. S6).

**FIG 2.**
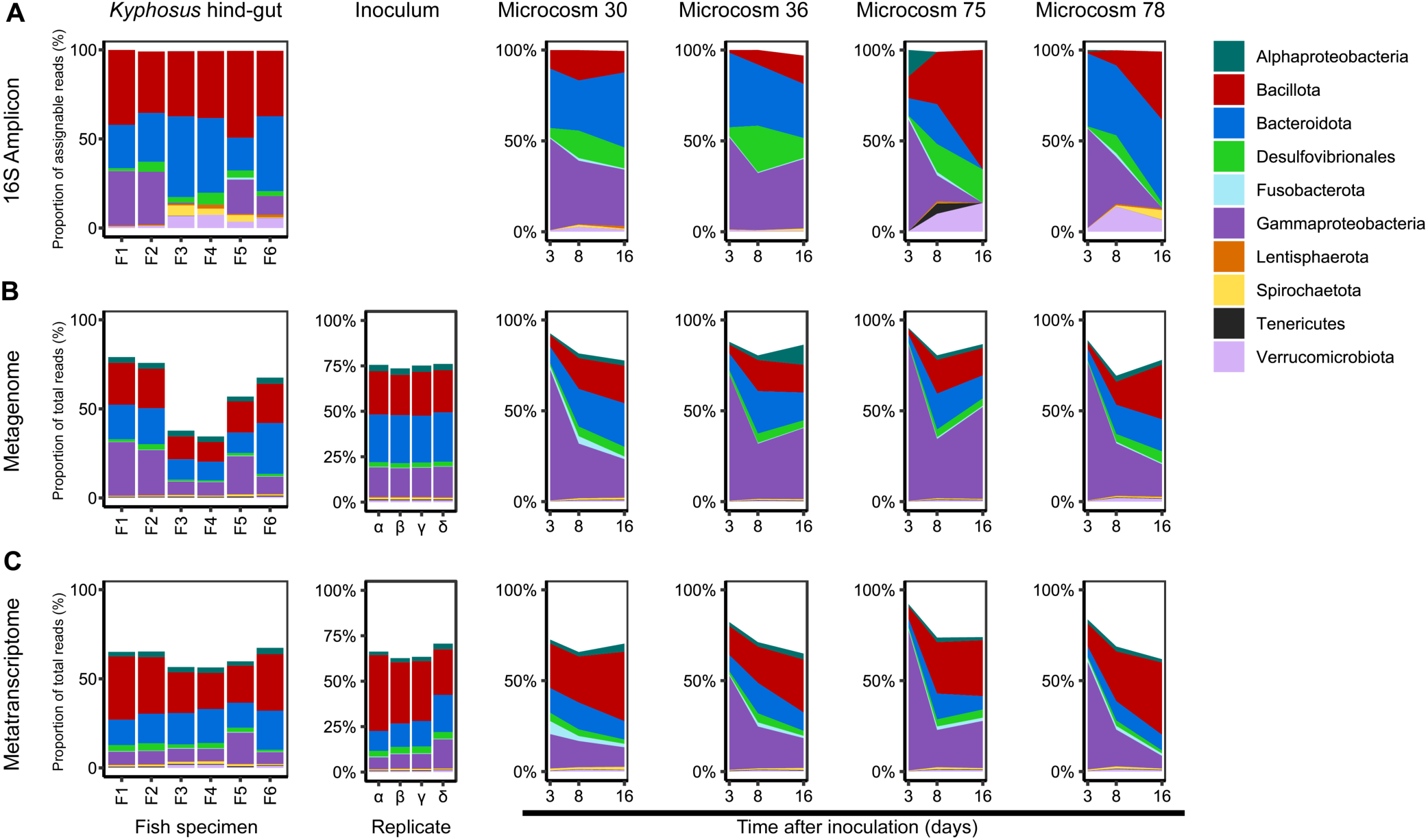
Overall composition and expression profile of microbial taxa from *K. cinerascens* natural hindgut, source inoculum, and selected *in vitro* microcosms. Taxonomic assignments of the microbial (A) 16S rRNA gene amplicon sequences, (B) metagenomic reads, and (C) metatranscriptomic reads from the microbial communities in 6 *K. cinerascens* natural hindgut samples, four replicates of the inoculum mixture used to prepare all the microcosms Three timepoints (3, 8 and 16 days after inoculation) are shown for the selected *in vitro* microcosms with the four culture conditions that propagated the largest number of microbial taxa from the active members of the natural hindgut samples.

The simultaneous extraction of DNA and RNA from the same biological material enabled the parallel comparison of the representation and transcriptomic profiles in the microbiomes from natural *K. cinerascens* hindgut and *in vitro* samples. To assess the expression of microbial enzymes that can be confidently associated with the biotransformation of large macroalgal polysaccharides, putative enzymes biotransforming green macroalgae polysaccharides (e.g., ulvan lyases) were excluded from this analysis because they could not be confidently connected to a single substrate (Bäumgen, Dutschei, and Bornscheuer 2021).

The initial 1,153 MAGs obtained were de-replicated into 471 populations, and a subset of 92 dereplicated high-quality MAGs (>95% completeness, <5% contamination) from lineages expressing transcripts for enzymes targeting macroalgae constituents (Fig. 3) were further assessed regarding their metabolic potential and temporal expression of enzymes for the biotransformation of sulfated polysaccharides and polyphenols (Fig. 4). Overall, *in vitro* propagation induced an increase of transcripts related to all the polysaccharide- and polyphenol-degrading genes assessed here, which generally plummeted sometime between eight and 16 days after inoculation (Fig. 3), potentially due to the transformation of their substrates into simpler molecules. The only exception was microcosm 30, which presented the simplest combination of macroalgae in its media among the selected microcosms. In this case, sometime between zero and three days post inoculation the overall expression of genes related to the degradation of alginate dropped to a third of their abundance compared to levels present in the initial inoculum (Fig. 3A). Furthermore, in microcosm 30, the expression of genes related to the degradation of carrageenan and fucoidan plummeted between three and eight days after its inoculation (Fig. 3A).

**FIG 3.**
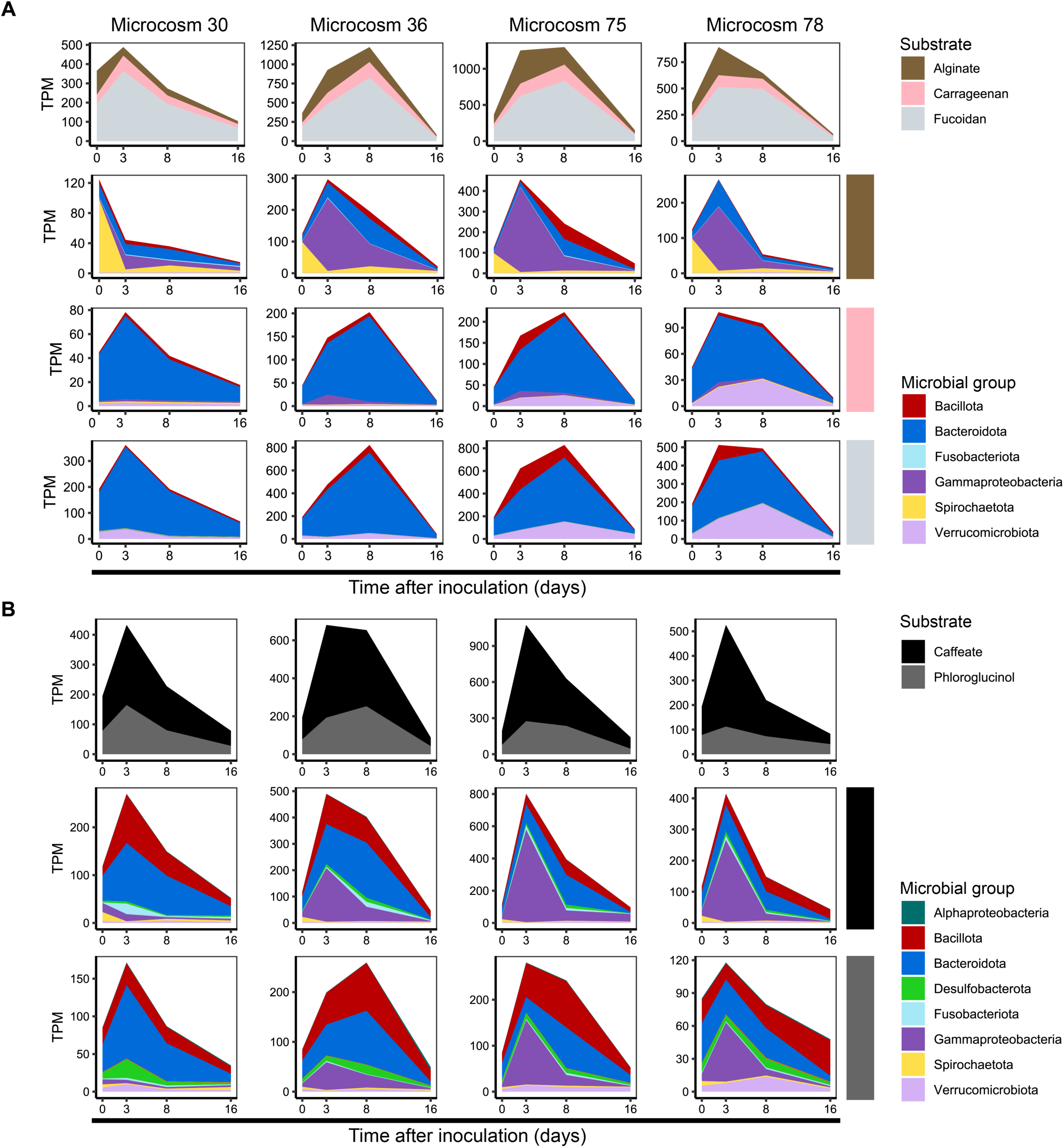
Expression of enzymes biotransforming macroalgae constituents

**FIG 4.**
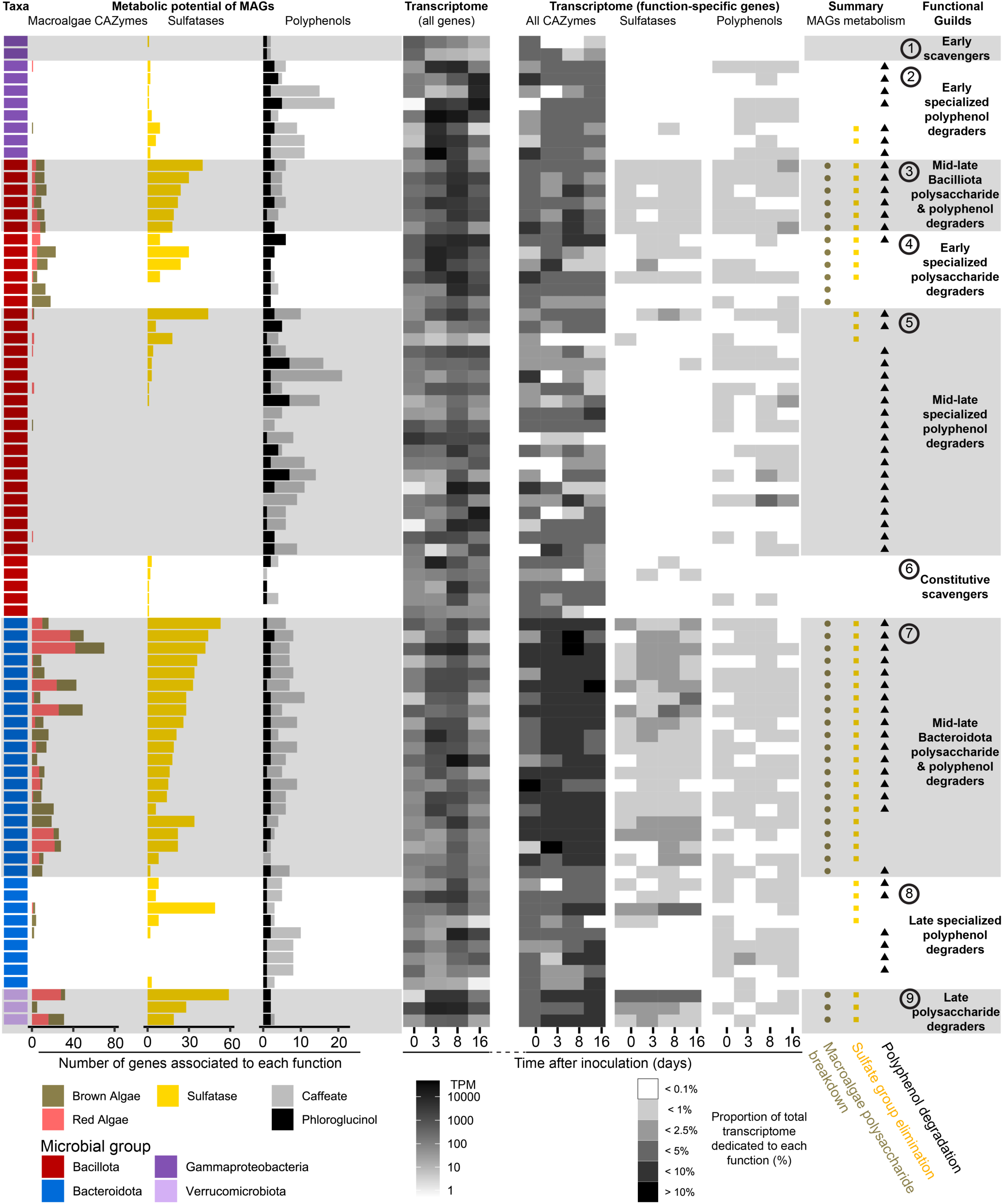
Functional guilds of the *K. cinerascens* MAGs and their transcriptomic profiles along a 16-day *in vitro* cultivation experiment. From left to right, the first column denotes the taxonomic assignment of the MAGs. The bar plots display the total gene count for CAZymes involved in the biotransformations of macroalgae polysaccharides, sulfatase genes, and genes involved in the breakdown of caffeate and phloroglucinol. Black and white heatmaps show the overall representation of these MAGs in the entire metatranscriptomes, then the recruitment of transcripts related to function-specific gene categories: all CAZymes, sulfatases, and genes involved in the biotransformation of caffeate and phloroglucinol. The two rightmost sets of columns provide metabolic summaries for each MAG, which were used to classify them into functional guilds within each taxon according to their capacities to biotransform and/or utilize macroalgal constituents. CAZyme categories representing the initial breakdown steps of alginate, xylan, carrageenan, agar, and fucoidan were used to differentiate between microbes that lyse large macroalgal polysaccharides and those that scavenge on smaller oligo-and monosaccharides.

When comparing the microbiomes of the selected microcosms, the proportion of Gammaproteobacteria transcripts in microcosm 30 that were detected three days after inoculation were less than half the representation of this group in the metagenome from the same time point (Fig. 3B and C). The reduced representation of Gammaproteobacteria transcripts was also evident when comparing microcosm 30 to the other selected microcosms regarding the expression of genes involved in the degradation of alginate (Fig. 3A) and polyphenol-related enzymes (Fig. 3B).

The family Vibrionaceae is the most abundant Gammaproteobacteria group propagated in all microcosms (Fig. 2 and Fig. S4). Among MAGs generated from the *K. cinerascens* natural hindgut and *in vitro* metagenomes, the Vibrionaceae populations with the highest representation were related to the genera *Photobacterium*, *Vibrio*, and *Shewanella* (Table S6).

### Identification of microbial guild structure in macroalgae processing

The overall transcriptional profiles of the Gammaproteobacteria MAGs revealed the presence of two functional guilds (Fig. 4). One guild expressed enzymes involved in polyphenol degradation during the early stages of microcosm incubation, particularly before day eight, likely representing specialized populations initiating the detoxification of macroalgal polyphenols. The other Gammaproteobacteria functional guild expressed CAZyme genes associated with the degradation of the relatively bioavailable polysaccharide alginate during this same early window. The temporal confinement of these activities to the initial phase of the experiment suggests that populations from the latter Gammaproteobacteria guild operated as opportunistic scavengers and early responders utilizing readily available substrates. Thus, the two *K. cinerascens* Gammaproteobacteria functional guilds detected here significantly decrease their *in vitro* metagenomic and metatranscriptomic representation because they are likely responsible for early polyphenol degradation steps and the utilization of less complex polysaccharides (Fig. 5). This is in agreement with previous studies showing a higher proportion of Vibrionaceae among intermediate gut sections compared to the more distal hindgut communities of *K. cinerascens* (Sparagon et al. 2022) and other *Kyphosus* fish (Pisaniello et al. 2023). These same studies have shown that the representation of other microbial phyla, which express the majority of the genes involved in polysaccharide breakdown and polyphenol degradation (*i.e*., Bacillota, Bacteroidota, and Verrucomicrobiota), increases in communities located in the more distal hindgut compared to those from intermediate gut sections of these herbivorous fish.

**FIG 5.**
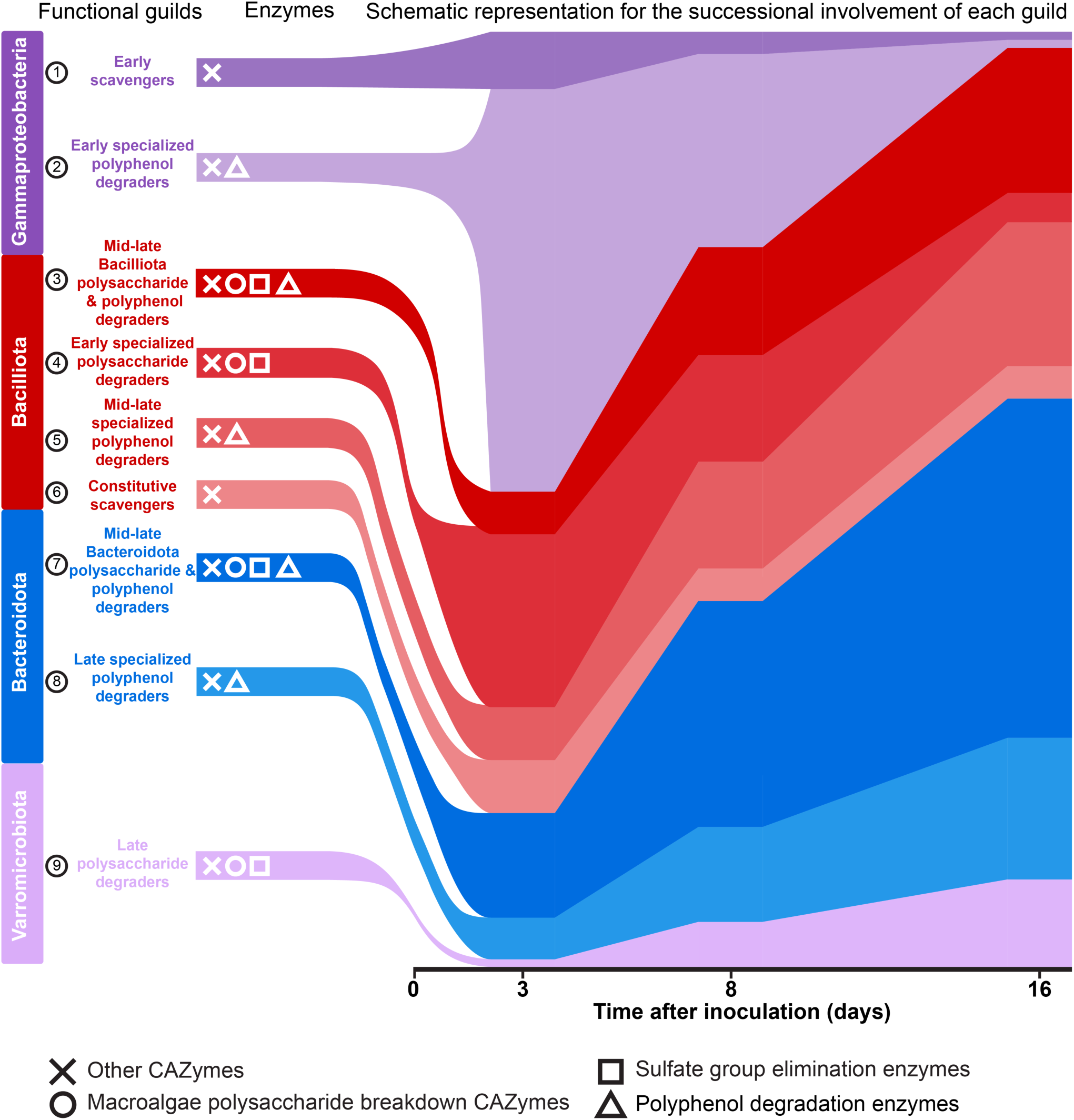
Transcriptional successional involvement of the main functional guilds involved in macroalgae biotransformations in the *K. cinerascens* 16-day *in vitro* cultivation experiments.

Transcripts assigned to the Bacillota phylum were involved in the processing of all polysaccharides and polyphenols irrespective of the media compositions assessed here (Fig. 3). Still, the extent of their involvement was proportionally more important regarding the expression of enzymes involved in polyphenol degradation (Fig. 3B). Bacillota MAGs assigned to the genera *Faecalibacterium*, *JAKSVV01*, and *Caccovivens* showed a broader range of functional capacities and temporal expression patterns (Fig. 4). Bacillota microbes presented four functional guilds, of which the first one included members capable of both polysaccharide and polyphenol degradation, with pronounced transcriptional activity at days eight and sixteen (Fig. 4), suggesting a role in processing more structurally complex or partially transformed substrates. A second guild of Bacillota MAGs appears specialized in polysaccharide breakdown and followed an earlier transcriptional activity pattern, with higher overall transcriptional levels at days three and eight (Fig. 4). The third set of Bacillota MAGs comprised a functional guild specialized in polyphenol degradation and had a higher transcriptional activity mostly at days eight and sixteen (Fig. 4). Finally, the fourth guild of Bacillota MAGs appeared to act as generalist scavengers, with no clear transcriptional pattern throughout the time series (Fig. 4). These trends suggest Bacillota include metabolically versatile taxa that are involved in distinct temporal niches within the successional dynamics of the macroalgal biotransformation process (Fig. 5).

Transcripts assigned to the Bacteroidota phylum were proportionally more important regarding the expression of enzymes for polysaccharide degradation (Fig. 3A), but were also one of the main microbial taxa involved in the processing of polyphenols irrespective of the media compositions assessed here (Fig. 3B). MAGs from the phylum Bacteroidota were predominantly associated with the genus *Alistipes* and presented high transcriptional levels at the intermediate (day eight) and final (day sixteen) timepoints. Previous studies have reported that hindgut *Alistipes* populations from other *Kyphosus* species have the potential to utilize a wide range of macroalgal polysaccharides, thanks to the high number of CAZyme gene clusters in their genomes (Facimoto et al. 2024); (Facimoto et al. 2025). Overall, Bacteroidota MAGs with both polysaccharide and polyphenol degradation capacities exhibited elevated transcriptional activity from days three to sixteen, indicating a role in the continued degradation of polysaccharides and breakdown of polysaccharide substrates throughout the *in vitro* experiments (Fig. 5). A second Bacteroidota functional guild, that seemed to specialize solely in the degradation of polyphenol, had higher overall transcriptional activity towards the final time points of the experiment (Fig. 5).

Microcosms supplemented with the most complex macroalgal mixtures exhibited higher levels of Verrucomicrobiota transcripts associated with the degradation of carrageenan and fucoidan (Fig. 3A), as well as the polyphenol phloroglucinol (Fig. 3B). MAGs from the phylum Verrucomicrobiota were mostly assigned to the genus *CAKUIA01* (order Opitutales) and a high-quality MAG from the order Kiritimatiellales. These populations were primarily specialized polysaccharide degraders, with gene expression profiles indicating peak activity at the middle and final timepoints of the *in vitro* experiments (Fig. 5). These findings suggest that Verrucomicrobiota are particularly responsive to complex, heterogeneous mixtures of macroalgal compounds, likely benefiting from the metabolic activities of early-degrading populations and contributing to the later stages of macromolecule deconstruction.

Together, these results support the existence of a stepwise, collaborative degradation process among distinct microbial populations within the *K. cinerascens* gut microbiota. Early-acting specialists rapidly initiate the detoxification and breakdown of accessible substrates, while later-acting populations respond to the progressive transformation of more recalcitrant compounds. This ecological succession underscores the metabolic interdependence of gut symbionts in processing complex macroalgal diets.

Community composition at the metagenomic level and community functional expression patterns of the *in vitro* microcosms (Fig. 3) resembled the abundance profiles of 16S rRNA amplicon (Sparagon et al. 2022)(Pisaniello et al. 2023) and community sequencing (Facimoto et al. 2024). These prior studies assessed the fine-scale transition observed along the natural gastrointestinal tract of herbivorous fishes, including the shift from stomach to hindgut compartments, where increasingly processed substrates support the activity of more specialized microbial taxa from the Bacteroidota, Bacillota, and Verrucomicrobiota phylum. This research further validates the ecological relevance of the *in vitro* microcosm system developed here, as well as the potential to explore additional factors relevant to the natural gut such as oxygen tolerance and microbiome stability over long timescales.

Previous metatranscriptomic studies, in the closely-related *Kyphosus sydneyanus* fish (Facimoto et al. 2024), have shown that the biotransformations of macroalgae polysaccharides in natural samples of the two most distal sections of the hindgut are mainly performed by the Bacteroidota (class Bacteroidia), and Bacillota (classes Clostridia and Bacilli). The findings of this study build on this foundational work by expanding the identification of microbial groups that are transcriptionally active members in natural hindgut microbiomes through cDNA-based 16S rRNA gene amplicon sequencing and metatranscriptomic approaches, and add a novel temporal dimension to the expression dynamics of anaerobic macroalgae-degrading microbiota using *in vitro* microcosms. These insights not only revealed when specific functional guilds are active but also how their interactions unfold over time in response to substrate availability, adding a new layer of resolution to the ecology of generalist herbivorous fish gut microbiomes. Furthermore, this is the first study assessing the biotransformation capacities of macroalgal polyphenols by complex *in vitro* microbiomes that closely resemble the natural hindgut microbiome of generalist herbivorous fish. However, because phloroglucinol is only one of the final breakdown products of diverse polyphenols, the degradation of macroalgae polyphenols likely involves many additional enzymes beyond those assessed here. As a result, our evaluation of polyphenol-related enzyme expression was constrained by the current range of annotated targets available in the CAMPER database (McGivern et al. 2024). Thus, future studies identifying enzymes with validated capacities to biotransform physiologically relevant macroalgae polyphenol substrates (Cotas et al. 2020) and identifying their gene sequence will be vital to developing much needed bioinformatic strategies to identify these enzymes.

## MATERIALS AND METHODS

### Fish sampling

*Kyphosus cinerascens* specimens were collected by local fishers using a spear gun directly offshore of the Ocean Era facility at Keahole Point, Kona, Hawai’i Island, USA on July 7th 2022 (19.7286, −156.0619). Fish were euthanized by pithing, in accordance with IACUC protocol S12219, and then bagged and maintained cold with frozen ice packs for about an hour until their GI was sampled for cultivation and nucleic acid extraction. Biometrics such as mass and fork length (the distance between the snout and the fork of the tail fin) were measured for each fish (Table S1). Specimens were dissected as described previously (Sparagon et al. 2022) to collect the microbiota from the distal end (hindgut) section of the host intestine under anaerobic conditions.

### Propagation of fish gut-microbiota

To generate media combinations that mimicked the dietary elements of the *Kyphosid* fish host, macroalgae *Turbinaria* spp., and *Dictyota* spp. were collected from Keahole Point (Kona, Hawai’i Island, USA; 19.7286, −156.0619), while *Ulva* spp., *Halidrys* spp., and assorted Coralline macroalgae were collected from La Jolla Tide Pools (San Diego, California, USA; 32.841566, −117.281807). The red *Agardhiella subulata* macroalgae was cultivated on land (Augyte et al. 2023) at the Ocean Era facilities (73-970 Makako Bay Dr, Kona Island, Hawaii, USA). Seawater previously passed through a 0.22 m pore size filter (Sterivex, Millipore) was used to soak and triple wash the macroalgae -to remove sand, epibionts, and macrofauna - before being blended and freeze dried into a fine powder. The Artificial Sea Water (ASW) of 35 practical salinity units used here consisted of Instant Ocean, 40 g/L (Spectrum Brands, Blacksburg, VA). Additionally a Basal Salts Medium (BSM) was formulated to replicate the natural physico-chemical parameters of marine bony fish gastrointestinal-tract (see “Anaerobic in vitro microcosms to propagate *K. cinerascens* gut microbiota”). Resazurin was added at 0.005 g/L final concentration to visually examine O_2_ levels of all cultures. A total of 4 g/L of freeze-dried macroalgae combinations were added to each culture (Fig. S1 and Table S2). The hindgut contents of six *K. cinerascens* fish were combined to serve as inoculum for all the microcosms of this study. The gut inoculum was mixed with anaerobic BSM and sieved through a sterile 1 mm strainer in the anaerobic chamber. Microcosms were inoculated with the sieved *K. cinerascens* hindgut content mixtures at a final 1:180 dilution ratio in 50 mL 100 mL serum bottles crimp-sealed with a rubber septum. The 50 mL of headspace was flushed with up to ∼29 PSI with N_2_:CO_2_ 80:20 gas. All cultures were maintained at 22 °C during the entire experiment. When sampling each timepoint, the exposed surface of the serum bottle crimper and rubber stopper were sterilized with alcohol pads before retrieving 1 mL of each culture. These samples were mixed with 1 mL of 2X DNA/RNA Shield solution (Zymo) and then stored at −80 °C till further processing. 150 L of each natural sample, culture, and the non-inoculated media controls were aliquoted into 96-well plates to screen their microbiomes.

#### Anaerobic *in vitro* microcosms to propagate *K. cinerascens* gut microbiota

Since the microbiota residing in the distal intestine section (hindgut) of *Kyphosus* fish have the highest proportion of genes involved in the biotransformation of macroalgae (Podell et al. 2023), lumen samples from this section were collected from six *K. cinerascens* specimens to serve as inoculum for the various microcosms of this study and offer a reference for the natural microbiome composition. The anaerobic *in vitro* microcosms generated here were designed using media combinations (Fig. S1; Table S2) that replicated the host dietary items (Wu et al. 2022; Choat, Clements, and Robbins 2002)(Sparagon et al. 2022). Additionally a Basal Salts Medium (BSM) was formulated to replicate the natural physico-chemical parameters of marine bony fish gastrointestinal-tract regarding the overall drop in Na^+^ plus Cl^-^ to the bloodstream through the GI-tract epithelium. This decreases the concentrations of these ions from >300 mM in sea water to 50-80 mM in the gastrointestinal-tract lumen while establishing high concentrations of Mg^2+^, SO_4_^2-^, and the saturation of carbonate (Marshall & Grosell 2006). Some microcosms were also amended with marine bony fish type II mucins (Jin et al. 2015), bile salts (Buchinger, Li, and Johnson 2014), and short-chain fatty acids (SCFAs) that have been previously detected in *Kyphosus* hindguts (Rimmer and Wiebe 1987; Pardesi et al. 2022) to favor the propagation of microbial groups that might require these nutrients (Oliver et al. 2024).

Specifically, BSM was a mixed solution of: MgSO_4_, 10.833g/L; K_2_CO_3_, 0.8293 g/L; CaCO_3_, 0.6 g/L; MgCO_3_, 1.6863 g/L; and a 1:10 dilution of ASW. Cysteine hydrochloride monohydrate (1028390100, Sigma) 0.5 g/L final concentration was added before autoclaving to the BSM to make it anaerobic. The following components were added to make different culture media combinations (Fig. S1 and Table S2): 3.62 g/L mucin Type II (M2378-100G, Sigma)(Jin et al. 2015); a final concentration of 0.226 g/L taurocholic acid sodium salt hydrate (T4009-5G, Sigma) and chenodeoxycholic acid (C9377-5G, Sigma) at a 4:1 ratio (Buchinger, Li, and Johnson 2014); Biotin, 2 mg/L (Sigma); folic acid, 2 mg/L (Sigma); pyridoxine acid, 10 mg/L (Sigma); riboflavin, 5 mg/L (Sigma); thiamine hydrochloride, 5 mg/L (Sigma); cyanocobalamin, 0.1 mg/L (Sigma); nicotinic acid, 5 mg/L (Sigma); P-aminobenzoic acid, 5 mg/L (Sigma); lipoic acid, 5 mg/L (Sigma); DL-pantothenic acid, 5 mg/L (Sigma).

### DNA ‘Fast-Extract’ and DNA-based 16S rRNA gene amplicon sequencing

A high-throughput and inexpensive “Fast-Extract” DNA extraction procedure was implemented here to process the hundreds of microcosms and the natural *K. cinerascens* hindgut samples. The titration of a mixed culture of *Bacillus subtilis* and *Paracoccus* spp. with known cell concentrations revealed that the absolute limit of detection for the Fast-Extract procedure coupled with 16S rRNA amplicon sequencing was between 31 and 16 cells (Fig. S1). Still, the number of reads assigned to *B. subtilis* and *Paracoccus* spp. fell within the same range as the background noise (*i.e*., quality filtered reads assigned to taxa that were not present in the initial mixed culture) when <1,600 cells were present in the biological sample (Fig. S1). Since >90% of the reads associated to the background noise were assigned to the Vibrionaceae family (*i.e*., from either the *Catenococcus* or *Vibrio* genus; Table S3) the observed sequence background noise is likely the result of well-to-well contamination (Minich et al. 2019) with neighboring microcosm samples processed for nucleic acid extraction in the same 96-well multiplates and sequenced parallely (Fig. 2).

For the Fast-Extract procedure, cell lysis of microcosm aliquots of 150 L were performed using a heat and freeze method. Specifically aliquots were heated to 95 °C for 10 min and then immediately placed at −20 °C for another 10 min. The samples were then placed on ice and thawed before being centrifuged at 2,000 *g* for 5 min at 4 °C. Immediately after centrifugation, 100 L of supernatant was transferred to a new 96-well plate to undergo a 1X bead cleanup step using AMPure XP beads (Beckman Coulter). Briefly, the magnetic beads were allowed to equilibrate to room temperature, and then vortexed 30 s at high speed before adding 1 volume (100 l) to each sample. The beads and samples were mixed by vortexing for 5 s at the highest speed and left to incubate at room temperature for 5 min. The plate was placed on a magnetic rack and the supernatant was discarded to remove cellular debris. While on the magnetic stand or plate, samples were washed twice with 2 volumes of freshly made ethanol 70 % (v/v) allowing for the beads to incubate for 30 s with this solution before removing it. Then the magnetic beads were air dried for 5 min at room temperature before removing them from the magnetic rack and gently resuspending them in 30 L of sterile DNase-free water. The samples were left to incubate for 10 min at 37 °C before the beads were decanted on a magnetic rack and the supernatant with the DNA was retrieved and stored at −20 °C for further processing. A variety of positive and negative controls were included in the DNA extraction steps. Positive controls were utilized to estimate the limit of detection of the assay and assess or ensure minimal contamination occurred following the Katharoseq protocol (Minich et al. 2018)(Minich et al. 2022). Seven serial dilutions of a *Paracoccus* spp. and *Bacillus subtilis* sp. mixed culture with an initial concentration of 1.6×10^9^ and 3.1×10^8^ cells/mL, respectively, were subjected to the same Fast-Extract procedure and 16S rRNA amplicon sequencing described above to compare input cell counts with the resulting sample composition to determine the absolute limit of detection of this procedure. Cell concentrations for the initial Paracoccus spp. and Bacillus subtilis sp. mixed culture was corroborated by flow cytometry as previously described (Minich et al. 2018). The V4 region (∼290 bp) of the 16S rRNA gene was amplified using a two-step PCR protocol to create dual-barcoded amplicons. The first reaction used primers 515F-Y and 806rb with overhangs for attachment of Illumina-compatible indexes in the second reaction (Caporaso et al. 2012). The initial reaction was performed in triplicate using a high-fidelity Q5 polymerase (NEB, Ipswitch, MA) with the following steps: initial denaturation of 30 s at 98 °C; 25 cycles of 10 s at 98 °C, 20 s at 50 °C, 30 s at 72 °C; final extension of 2 min at 72 °C. The second reaction was performed as described above except the amplification steps consisted of only 8 cycles with an annealing temperature of 56 °C. Barcoded amplicons were cleaned using AMPure XP beads (Beckman Coulter, Brea, CA), pooled at equimolar concentrations, and sequenced using Sp500 chemistry (2×250) on a MiSeq instrument (Illumina, San Diego, USA).

### Parallel DNA and RNA extractions and generation of cDNA

A 2 mL sample of the natural hindgut contents of each *Kyphosus* fish, as well as composite mixtures of all six fish hindgut samples (used to inoculate cultures) were preserved by mixing in 3 volumes of DNA/RNA Shield solution (Zymo), then stored at −80 °C until further processing. These natural samples and a select number of cultures were processed for simultaneous parallel extraction of nucleic acids from the same biomaterial using the DNA and RNA Purification protocol of the ZymoBIOMICS MagBead DNA/RNA kit (Zymo). Samples were processed according to the manufacturer’s instructions except that 750 L of biomaterial was used in each extraction, with the volume of new DNA/RNA Shield solution added initially to each bead beating tube reduced to 350 L. Other modifications to the manufacturer’s recommended procedure were that all samples were bead-beaten for 40 min at 4 °C using a VortexGenie2, and the incubation with Proteinase K was skipped. Immediately after extraction, 11 L of RNA solution from the natural hindgut contents of each *Kyphosus* fish and four replicate samples of the inoculum mixture of all six fish hindgut contents was retro-transcribed into cDNA with the SuperScript III kit (Invitrogen) following the random primer procedure.

### 16S rRNA gene amplicon sequence analysis

All amplicon sequence data were analyzed together using the DADA2 package (v1.11.3)(Callahan et al. 2016) implemented in R (v3.4.4; Tierney 2012). The primers were removed using CutAdapt (v1.2.1; Martin 2011), and the sequences from each pair were trimmed and quality filtered (maxN = 0, maxEE=c(2,2), truncQ = 2, rm.phix=TRUE, trimLeft=10, trimRight=20, minLen=160). Amplicon Sequence Variants (ASVs) were inferred from de-replicated sequences. Taxonomic assignment was performed using the Genome Taxonomy Database (GTDB v202) for Bacteria and Archaea (Chaumeil et al. 2019). Chimeras were removed with ‘removeBimeraDenovo’ using the “consensus” removal method. All sequences assigned to mitochondria or chloroplasts were removed, and DECONTAM (Davis et al. 2018) was used to further identify contaminants within these datasets using the “frequency” method with a default probability threshold (TRUE if *p* < 0.1) to discriminate foreign taxa observed in the negative controls of autoclaved media constituents without hindgut inoculum (*i.e*., microcosms no. 1 to 15). Samples from these negative controls and those with less than 1000 reads were excluded leaving a total of 23 natural hindgut samples (11 from the DNA and 12 from the cDNA fractions) and 172 microcosm samples. Differences of taxonomic abundance between DNA and cDNA samples were calculated using Tukey’s honestly significant different test at p < 0.05 (Abdi and Williams 2010). The proportion of ASVs in each sample was plotted with the Phyloseq package (McMurdie and Holmes 2013). Alpha diversity was evaluated using Breakaway (Willis and Bunge 2015) with unaltered frequency counts of quality-filtered and decontaminated ASVs.

We used the ASV count table to construct a network of samples (ANET-samples), applying the Association Network (Anets) framework to assess associations between pairs of samples based on the similarity of their ASV co-occurrence patterns (Karpinets et al. 2018). Briefly, the OTU table was transformed into a matrix where both rows and columns represent samples, and each cell contains the number of shared ASVs between each sample pair. To quantify similarity, we calculated the Pearson correlation coefficient for each pair of samples. Pairs with a correlation coefficient (R) ≥ 0.70 were considered associated. The resulting correlation matrix was converted into a Cytoscape-compatible dataset using R, and sample networks were visualized in Cytoscape v3.6.1 (https://cytoscape.org; Shannon et al. 2003; Ono et al. 2025). Modularity of the sample network was assessed using Markov Cluster (MCL) assignments based on correlation thresholds of R ≥ 0.70. In addition, ASVs interaction networks were inferred using FlashWeave with the “sensitive” option enabled (Tackmann, Matias Rodrigues, and von Mering 2019). FlashWeave applies a centered log-ratio transformation to account for compositional data characteristics. Modularity in ASV networks was also computed via MCL clustering, using the same correlation threshold (R ≥ 0.70) to define modules. All statistical analyses and visualizations of the amplicon sequencing data were conducted using the Microeco package (v0.20.0; Liu et al. 2021) in R version 4.5.0.

### Metagenomic and metatranscriptomic sequencing

Barcode-indexed sequencing libraries were generated from the metagenomic DNA samples using the DNA KAPA Hyper Prep Kit (Kapa Biosystems-Roche, Basel, Switzerland) following the Earth Microbiome Project protocol (Sanders et al. 2019). The pools were quantified by qPCR with a Kapa Library Quant kit (Kapa Biosystems-Roche, Basel, Switzerland) and each pool was sequenced using 2×150 chemistry on an NovaSeq instrument (Illumina, San Diego, USA). Total RNA (250 ng) was subjected to the depletion of ribosomal sequences using the Qiagen QIAseq FastSelect kit following the manufacturer’s instructions (Qiagen, QIAseq FastSelect –rRNA Fish Kit combined with the 5S/16S/23S rRNA Kit–). The ribo-depleted RNAs were then used to generate strand-specific and barcode indexed RNA-seq libraries using the KAPA RNA HyperPrep Kit (Kapa Biosystems-Roche, Basel, Switzerland) following the instructions of the manufacturer. The fragment size distribution of the RNA libraries were verified via micro-capillary gel electrophoresis on a Bioanalyzer 2100 (Agilent, Santa Clara, USA), quantified by fluorometry on a Qubit fluorometer (LifeTechnologies, Carlsbad, USA) and pooled in equimolar ratios that were sequenced using 2×150 chemistry on an NovaSeq instrument (Illumina, San Diego, USA).

### Metagenomic and metatranscriptomic assembly and bioinformatic processing

Metagenomic reads were cleaned with fastp v.0.23.4 (Chen et al. 2018) with --trim_poly_g, and assembled with metaSPAdes v.3.15.5 (Nurk et al. 2017) and annotated with Prokka v.1.14.6 (Seemann 2014). Metagenomic binning was performed with COMEBin v.1.0.3 (Wang et al. 2024). MAG quality was assessed with CheckM2 v.1.0.2 (Chklovski et al. 2023) and taxonomic lineage was assessed with GTDB-Tk v.2.3.2 (Chklovski et al. 2023; Chaumeil et al. 2022) using Genome Taxonomy Database release 214 (Parks et al. 2021). MAGs were dereplicated with dRep v.3.4.2 using default parameters (Olm et al. 2017). A phylogenetic tree of MAGs was generated using PhyloPhlan v.3.1.68 (Asnicar et al. 2020) and visualized with the ggtree v.3.4.0 package (Yu et al. 2017) in the R v.3.6.1 environment. The taxonomic distribution of metagenomic and metatranscriptomic reads was determined using Kraken v.2.1.3 (Wood, Lu, and Langmead 2019) with a custom database utilizing all sequences in NCBI nr (Sayers et al. 2022) as of August 2022. Kraken outputs were converted to biom data files using kraken-biom (https://github.com/smdabdoub/kraken-biom) and then frequency and taxonomy tables were generated with ‘qiime tools extract’ in the Qiime2 environment (Bolyen et al. 2019). The metagenomic read-based community analyses were then performed using the same procedures for amplicon frequency and taxonomy data tables as described above.

Metatranscriptomic reads were cleaned with fastp v.0.23.4 (Chen et al. 2018) with the settings -- trim_poly_g -f 10 -t 2. All rRNA sequences that remained were computationally removed using KneadData v.0.10 (Beghini et al. 2021). Individual de novo metatranscriptomes were assembled using rnaSPAdes v.3.15.5 (Bushmanova et al. 2019) and transcripts of interest were selected using TransDecoder v.5.7.1 (https://github.com/TransDecoder/TransDecoder).

### Gene annotation and expression

Sequences associated with CAZyme genes were annotated using dbCAN v.4.1.4 (J. Zheng et al. 2023) based on results from DIAMOND (J. Zheng et al. 2023; Buchfink, Xie, and Huson 2015) and HMMER (Johnson, Eddy, and Portugaly 2010). Similarly, sulfatase gene sequences were annotated using hidden markov models provided by the SulfAtlas v.2.3.1 database (Stam et al. 2023), and genes for enzymes targeting polyphenols were annotated using CAMPER v.1.0.0 (McGivern et al. 2024).

The taxonomic assignment of the metagenomic and metatranscriptomic reads was done using Kraken v.2.1.3 (Wood, Lu, and Langmead 2019) with a custom NCBI nr v2022.08 database (Sayers et al. 2022). Gene expression values in transcripts per million (TPM) were calculated using RSEM v.1.3.3 (Li and Dewey 2011) by mapping them to MAGs assembled in this study using Bowtie v.2.5.4 (Langmead et al. 2019).

## ACKNOWLEDGMENTS

This publication includes data generated at the DNA Technologies and Expression Analysis Core at the UC Davis Genome Center, utilizing Illumina NovaSeq library preparation and sequencing instrumentation that was purchased with funding from a NIH Shared Instrumentation Grant 1S10OD010786-01. Computational analyses were performed using the San Diego Supercomputer Center’s Triton Shared Computing Cluster. This work was funded by the United States Government, Department of Energy, Advanced Research Projects Agency—Energy grant ARPA-E DE-FOA-0001858 to R.S.N. and L.M.L.L. with contracts to C.E.N., L.W.K., and E.E.A., National Science Foundation grants OCE-1837116 and EF-2025217 to E.E.A., and National Institutes of Health NIEHS grant R01-ES030316 to E.E.A.

## AUTHOR CONTRIBUTIONS

Alvaro M. Plominsky, Formal analysis, Investigation, Methodology, Project administration, Software, Validation, Visualization, Writing – original draft, Writing – review and editing | Aaron Oliver, Formal analysis, Investigation, Methodology, Project administration, Software, Validation, Visualization, Writing – original draft, Writing – review and editing | Carlos Henriquez-Castillo, Formal analysis, Investigation, Methodology, Visualization, Writing – original draft, Writing – review and editing | Sheila Podell, Conceptualization, Investigation, Methodology, Resources, Supervision, Writing – review and editing | Jeremiah J. Minich, Formal analysis, Investigation, Methodology, Writing – review and editing | Simona Augyte, Investigation, , Writing – review and editing | Jennica Lowell-Hawkins, Investigation, , Writing – review and editing | Neil A. Sims, Conceptualization, Project administration, Supervision, Writing – review and editing | Eric E. Allen, Conceptualization, Funding acquisition, Project administration, Supervision, Writing – review and editing

## DATA AVAILABILITY

Raw and processed (meta)genomic data and all the processing parameters will be made publicly available through NCBI SRA upon manuscript acceptance.

## ADDITIONAL FILES

### SUPPLEMENTAL MATERIAL

Supplemental Tables S1-S7 are available online only.

### SUPPLEMENTARY FIGURES

**FIG S1** Experimental design of *K. cinerascens* hindgut microcosms and their validation. The proportion of each macroalga used in the culture media is shown in parentheses. Vitamin concentrations are detailed in the Materials and Methods section. Marine fish bile salts were used at a final concentration of 0.226 g/L. The hindgut inoculum final dilution ratio corresponds to the wet weight dilution factor of the natural sample mix in each microcosm. A “microcosm identifier number” was given to both negative controls that contained only sterilized components (microcosms number 1-15) and experimental cultures that were inoculated with the same hindgut inoculum (microcosms number 16-80).

**FIG S2** Fast-Extract limit of detection for 16S rRNA amplicon sequencing. Quality-filtered reads assigned to *Paracoccus* spp., *B. subtilis* sp. throughout a dilution series with known cell numbers are compared to those assigned to taxa that were not present in the initial biological material processed through this DNA extraction method.

**FIG S3** Breakaway diversity estimate for the *Kyphosus cinerascens* natural hindgut microbiome and its propagated *in vitro* communities. Source DNA (n=11) and cDNA (n=12) samples are shown after incubation at day 3 (T03, n=62), 8 (T08, n=59), and 16 (T16, n=52) are shown. Diversity estimates are based on all the ASVs present in each of the samples after quality filtering and removal of contaminants (mitochondria, chloroplasts, and contaminant ASVs detected in negative controls of autoclaved media constituents without hindgut inoculum) and artificial sequences (*i.e*., chimeras).

**FIG S4** Composition of the *Kyphosus cinerascens* natural hindgut microbiome, at the family level, showing microcosms that propagated the largest number of microbial families. The upper and lower schematics depict the presence/absence and proportional abundance, respectively, of microbial families comprising >5×10^-4^ % of the total ASVs in at least one sample of the natural *K. cinerascens* hindgut active microbiome (consisting of up to two replicates ⎼“rep_1” and “rep_2”⎼ per sampled specimen which were numbered from 1 to 6; Table S1) either within the DNA (n=11) or cDNA fractions (n=12). Microbial community composition is shown for microcosm media composition numbers 30, 35, 75, and 78 (Fig. S1 and Table S2) sampled at days 3 (T03), 8 (T08), and 16 (T16). The graph at the bottom of the figure represents the proportion of each taxon, considering all ASVs that were taxonomically assignable at the family level. “NA” denotes a time point that was discarded because it had <1,000 total sequences assigned at the family level.

**FIG S5** ASVs differentially represented in the *Kyphosus cinerascens* natural hindgut microbiome and the communities propagated through its microcosms. (A) The relative abundance of each of the top 28 ASV modules (Table S7) within each sample as determined through a correlation analysis network, grouped using Markov Cluster Algorithm (MCL). Modules comprise ASVs present in at least 5 samples and present a significant (corr > 0.7) co-association. The network was transformed to consider that edges are all positive. (B) Redundancy Analysis (RDA) to show how the media components (black arrows) explain the observed variation in the samples (dots). Colors denote the samples with regards to each MCL cluster. Ellipsoids are defined by a 95% confidence interval. The clustering of microcosm #78 at eight and sixteen days after inoculation are denoted in both sections of the figure to showcase their similarity to cDNA and DNA microbiota profiles from natural fish samples grouped within clusters one, two, and three.

**FIG S6** Phylogeny and biological origin for the initial (non-dereplicated) MAGs generated in this study for the main microbial groups comprising the *K. cinerascens* hindgut microbiome.

